# Temporal Dynamics of Antigen-Specific T Cell Expansion in Primary SARS-CoV-2 Infection

**DOI:** 10.1101/2025.11.04.686558

**Authors:** Andrew G. T. Pyo, Adlina Chan, Joshua Rosenheim, Clare Thakker, Lucy Bell, Gayathri Nageswaran, Suzanne Byrne, Ben Killingley, David Scobie, Mariya Kalinova, Alison Boyers, Andrew Catchpole, Alex Mann, Rik G. H. Lindeboom, Marko Z. Nikolic, Sarah A. Teichmann, Helen Wagstaffe, Mahdad Noursadeghi, Christopher Chiu, Curtis G. Callan, Andreas Tiffeau-Mayer, Ned S. Wingreen, Benny Chain

**Affiliations:** Department of Physics, Princeton University, Princeton NJ, USA; Department of Applied Physics, Stanford University, Stanford CA, USA; Division of Infection and Immunity, University College London, London, UK; hVIVO, London, UK; Wellcome Sanger Institute, Wellcome Genome Campus, Cambridge, UK; The Netherlands Cancer Institute, Amsterdam, NL; UCL Respiratory, Division of Medicine, University College London, London, UK; Cambridge Stem Cell Institute, University of Cambridge, Cambridge, UK; Department of Infectious Disease, Imperial College London, London, UK; Département de Physique, École Normale Supérieure, Paris, France; Institute for the Physics of Living Systems, University College London, London, UK; Lewis-Sigler Institute for Integrative Genomics, Princeton University, Princeton NJ, USA; Department of Molecular Biology, Princeton University, Princeton NJ, USA; Department of Computer Science, University College London, London, UK

## Abstract

Quantifying T cell response during primary infection in humans is crucial for understanding adaptive immunity. Leveraging a controlled human challenge to SARS-CoV-2, we characterized antigen-specific T cell response within and across individuals. Notably, individual clones reached similar maximum frequencies despite differences in the timing of their peak expansion. Mathematical modeling showed that this observation is consistent with precursor frequency, but not TCR signal strength, as the source of inter-clonal variability. Single-cell profiling revealed distinct temporal programs for CD4^+^ and CD8^+^ T cells, with CD4^+^ cells expanding earlier but contracting to a lower frequency. Clones with similar receptors, likely recognizing the same antigen, expanded at similar times. Together, these findings highlight how clone-intrinsic properties such as precursor frequency and lineage shape T cell clonal kinetics. These insights provide a quantitative framework for understanding T cell response in humans, with implications for vaccine design.

## Introduction

Following infection, antigen-specific T cells undergo clonal expansion during which they proliferate and differentiate to orchestrate and execute the immune response. This response is inherently clonal: each T cell expresses a unique T cell receptor (TCR), and those clones specific to the presented antigens activate and expand (*1, 2*). The kinetics of clonal expansion—how rapidly a clone expands, when it peaks and at what magnitude, and how quickly it contracts—are shaped by multiple factors, including TCR affinity, costimulatory signals, the initial precursor frequency of antigen-specific naïve T cells, and differentiation into long-term memory cells (*3*–*7*). While individual T cell repertoires vary substantially across individuals, population-level T cell responses to viral infection are remarkably reproducible (*8, 9*), suggesting underlying organizational principles that govern the immune response.

Insights into these principles have come largely from *in vitro* systems and *in vivo* murine models (*4, 5, 10*), which have elucidated how individual factors contribute to clonal dynamics. However, how these factors interact to shape the collective T cell response during primary infection in humans is less well understood. This gap is particularly important given the central role of clonal kinetics in determining the strength and quality of protective immunity, with broad implications for vaccine design, disease progression, and memory formation.

Studying T cell dynamics during natural infections in humans is inherently challenging, as key variables such as the timing, dose, and site of infection are typically unknown. To overcome these limitations, we utilized a controlled human infection study in which individuals were intranasally inoculated with a fixed dose of pre-Alpha strain of SARS-CoV-2 (*11*). This setting provides precise control over infection parameters and enables synchronized monitoring of both viral load and immune responses across individuals at unprecedented temporal resolution.

In the current study, we combined daily longitudinal TCR sequencing with mechanistic modeling to dissect the clonal dynamics of antigen-specific T cells during primary SARS-CoV-2 infection. We tracked individual T cell clones across 10 individuals, identified expanding clones using a statistical filtering approach, and quantified their proliferation and contraction kinetics. We then developed a mathematical model of clonal dynamics that incorporates variation in precursor frequency, TCR affinity, and viral load to interpret the observed kinetic heterogeneity across clones. To further examine how T cell lineage influences clonal behavior, we cross-referenced TCR sequences with single-cell transcriptomic data from the same cohort (*12*) to compare the dynamics of CD4^+^ and CD8^+^ T cell subsets.

Our findings provide a quantitative framework for understanding how variability in clonal kinetics arises in human T cell responses, shaped by both cell-intrinsic properties and by the anatomical and temporal context of infection. By disentangling these contributions, our work lays the foundation for future models of human adaptive immunity.

## Results

### SARS-CoV-2 controlled human infection model

The details of the study have been published previously (*11*). Briefly, 36 healthy adults aged 18 to 30 years were recruited to this single-center, phase 1, open-label, first-in-human study. Participants had no previous SARS-CoV-2 vaccination or documented infection and were excluded if positive for SARS-CoV-2 anti-S antibodies measured at screening. Two participants seroconverted between recruitment and challenge, and were excluded leaving 34 individuals. Participants were inoculated intranasally by pipette with 10 tissue culture infectious dose (TCID) 50 of a Good Manufacturing Practice wild-type SARS-CoV-2 strain (20A clade of the B.1 lineage having the D614G mutation; GenBank accession number OM294022) divided between both nostrils (100 μL each). Viral titers were measured at 12-hour intervals in mid-turbinate and throat swabs.

Strikingly, 17 out of 34 participants showed sustained viral replication which rose rapidly in the first few days post-inoculation. Viral load remained detectable for variable periods but was undetectable in all participants by day 28. Most of these individuals showed mild symptoms, but none became seriously unwell. The other 17 participants showed no sustained detectable viral replication, and are not analyzed further in this study.

### Characterization of expanded T cell clonal dynamics

To identify antigen-specific T cell clones from longitudinal clonal abundance measurements, we applied a filtering procedure that selected clones with low abundance at early timepoints (days –1 and 1 relative to inoculation) and statistically significant fold-increases between days 7–12 (see Materials and Methods). Expanding clones were identified using longitudinal abundance data for both α- and β-chain sequences in each individual. While the number of expanding clones varied by over an order of magnitude across individuals, the counts identified from α-chain and β-chain sequences were consistent within each individual (SI Fig. 1a). Of the 17 participants positive for viral replication, we included for subsequent analysis only those 10 patients for whom at least 10 expanding T cell clones were identified to enable analysis of relative timing and peak magnitude of responding clones. In total, we identified 502 expanding clones from α-chain data and 491 from β-chain data across these 10 individuals.

The longitudinal abundance measurements of clones identified as responsive to SARS-CoV-2 inoculation were then fitted to an analytical ansatz:

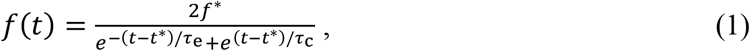

which describes exponential growth before the peak time *t*^∗^, where clonal dynamics transitions from expansion with timescale *τ*_e_, crossing over to contraction with timescale *τ*_c_ after the peak. The parameter *f*^∗^ = *f*(*t*^∗^) corresponds to the clonal abundance at peak time, where the clone transitions from expansion to contraction. The clonal kinetics for each expanding clone were fit to Eq. 1 (Fig. S1b,c). The aligned trajectories of expanding clones from all individuals—normalized by peak abundance and aligned by peak time—are shown in Fig. 1a,b, illustrating a consistent pattern of exponential expansion followed by exponential contraction.

**Figure 1.**
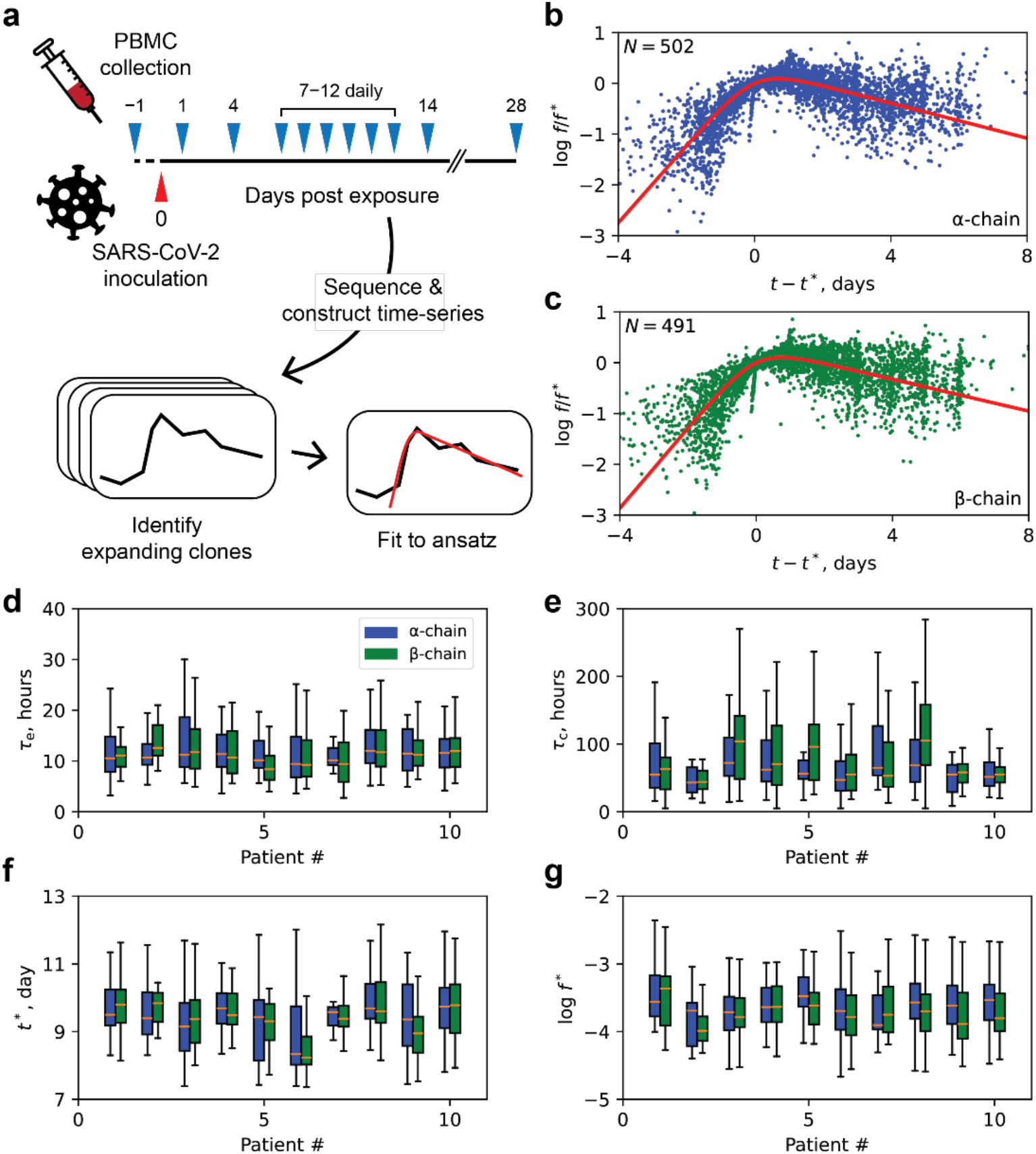
Characterization of T cell clonal dynamics in response to SARS-CoV-2. **a**, Schematic overview of the study design and the approach used to characterize T cell clonal dynamics. **b, c**, Abundance of expanding T cell clones inferred from CDR3 sequences of the α-chain **(b)** and β-chain **(c)** as a function of time relative to the peak time, estimated by fitting to Eq. 1. The average fit is shown in red. **d–g**, Inferred values of parameters characterizing T cell clonal dynamics. Expansion timescale **(d)**, contraction timescale **(e)**, peak day **(f)**, and peak abundance **(g)** of expanding T cell clones, from model fits for 10 individuals. Expanding T cell clones inferred from CDR3α sequences are shown in blue, and expanding clones inferred from CDR3β sequences are shown in green.

To quantify the kinetics of antigen-specific T cell clones, we compared the parameters obtained from fitting their trajectories to Eq. 1. As shown in Fig. 1d–g, we observed consistent parameter values between α-chain and β-chain sequences within individuals, as well as similar distributions across different individuals. In particular, we found a consistent expansion timescale of approximately 12 hours (equivalent to a doubling-time of 8 hours) (Fig. 1d), followed by a contraction timescale of approximately 3–4 days (equivalent to a half-life of 2–3 days) (Fig. 1e), though the contraction rate exhibited greater variability both within and across individuals. The average peak time of expanding clones was consistent across individuals, occurring at 9.5 days post-inoculation (Fig. 1f), and the average peak abundance reached approximately 1 in 4,000 T cells (Fig. 1g), with the largest clones reaching frequencies as high as 1 in 200 T cells. A summary of the inferred kinetic parameters is provided in Table 1.

**Table 1.** Inferred parameters from fits to clonal abundance time-series. Parameters were estimated by fitting the time-series of clonal abundance to Eq. 1. Errors indicate the standard deviation across 10 individuals.

| | CDR3α | CDR3β |
| --- | --- | --- |
| $\tau_e$ | $12.7 \pm 1.0$ hours | $12.3 \pm 1.6$ hours |
| $\tau_c$ | $80 \pm 13$ hours | $93 \pm 26$ hours |
| $t^*$ | $9.5 \pm 0.3$ days | $9.5 \pm 0.4$ days |
| $\log f^*$ | $-3.6 \pm 0.1$ | $-3.7 \pm 0.1$ |

### Peak abundance and peak day of antigen-specific T cell clones are uncorrelated

Following the quantification of clonal kinetics, we sought to determine the hierarchy of factors that determine the dynamics of antigen-specific T cells at the population level during primary infection. Specifically, is clonal behavior primarily driven by antigen stimulation strength (a combination of antigen dose and TCR affinity), or by precursor frequency? These two factors are known to influence different aspects of the T cell response; antigen signal is thought to regulate the proliferative capacity of a clone (*4*), while precursor frequency affects the likelihood and hence timing of recruitment into the response, as well the perceived burst size. Accordingly, if clonal expansion were primarily driven by one of these factors, we would expect a specific, distinct relationship between the timing and magnitude of expansion. For example, if precursor frequency were the main determinant, larger clones might exhibit earlier peak days due to having a higher chance of antigen encounter. Conversely, if affinity were dominant, clones with higher peak abundances might arise later due to the increased proliferative capacity.

To assess these possibilities, we examined the relationship between the peak day and peak abundance of each expanding T cell clone. Although precursor frequencies are typically below the detection threshold, our longitudinal sampling allowed us to capture clonal dynamics once they rise above this threshold—providing a window into the transition from the expansion to the contraction phase. As shown in Fig. 2a, we observed no apparent correlation between peak day and peak abundance across all 10 individuals, despite a 4–5 day spread in the timing of peak responses. To test this more rigorously, we computed the Spearman rank correlation coefficient for each individual; none showed a statistically significant association between the timing and magnitude of expansion (Fig. 2b). This lack of correlation suggests that the timing of clonal activation and the extent of clonal proliferation are governed by distinct and decoupled mechanisms. What, then, does this imply about the underlying drivers of clonal dynamics? To address this, we turned to mathematical modeling to explore how precursor frequency and affinity might influence the kinetics of clonal expansion.

**Figure 2.**
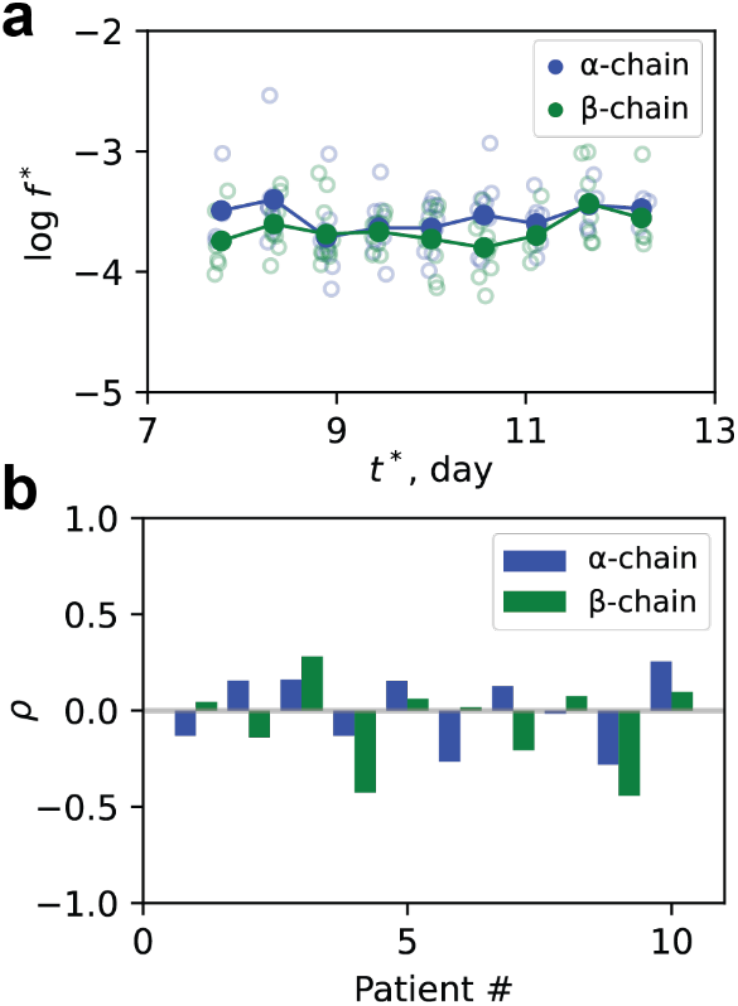
Peak abundance and peak day of antigen-specific T cell clones. **a**, Scatter plot of log peak abundance binned with respect to the peak day, as estimated from fits to Eq. 1 for each expanding T cell clone. Empty circles indicate the binned average within an individual, and filled circles indicate averages across individuals (with averages taken of logs). **b**, Spearman rank correlation coefficient between peak abundance and peak day computed separately for each individual.

### Mathematical model of T cell clonal dynamics

To investigate how precursor frequency and TCR affinity shape T cell clonal dynamics, we developed a hybrid stochastic-deterministic model of T cell activation and expansion (see Fig. 3a for model schematic). In this framework, naïve T cells interact with antigen-presenting cells (APCs), forming transient immune synapses that can either dissociate or lead to T cell activation. Given that T cell activation is a rare event during primary infection due to the paucity of antigen-specific T cells (*6*), we model activation as a stochastic process, where naïve T cells in clone *i* become activated at a rate dependent on viral load *r*_*i*_(*V*) (see Supplemental Information for derivation):

**Figure 3.**
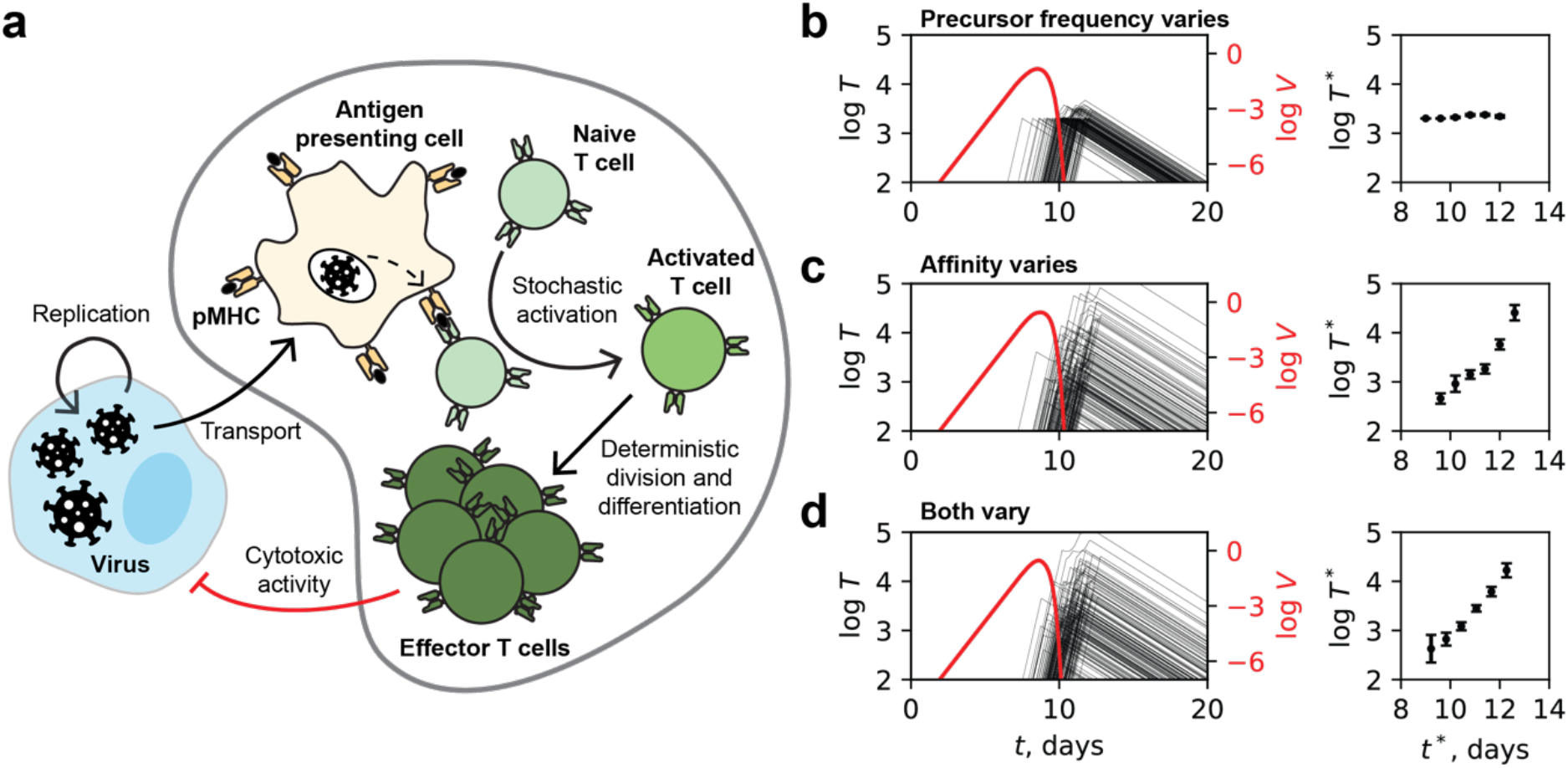
Mathematical model of T cell clonal dynamics. **a**, Schematic of the processes included in the model. **b–c**, Simulated trajectories of antigen-specific T cell clones (black) and viral load (red) are shown in the left panels; corresponding peak T cell abundances, binned by peak day, are shown in the right panels. We consider three scenarios: **(b)** clones vary in precursor frequency, modeled as log *N*_*i*_ = 1 + 0.2*ξ*_*i*_, where *i* indexes the clone and *ξ*_*i*_ ∼ *𝒩*(0,1) is a Gaussian random variable with zero mean and unit variance; **(c)** clones vary in affinity, with 1/*K*_*i*_ = 10^−5^(1 + 0.2*ζ*_*i*_) where *ζ*_*i*_ ∼ *𝒩*(0,1); and **(d)** clones vary in both precursor frequency and affinity. Error bars indicate standard error. See Supplementary Information for model details and parameter values.

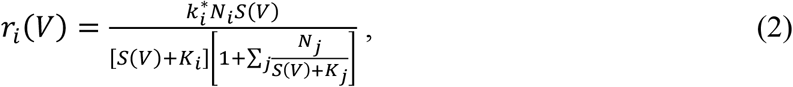

where 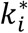is the maximum activation rate of each naïve T cell in clone *i, N*_*i*_ is the number of naïve T cells in clone *i, S*(*V*) denotes the effective amount of antigen presented on APCs as a function of viral load *V*, and 1/*K*_*i*_ is the effective affinity of clone *i*. Eq. 2 is derived under the assumption that at any time only a small fraction of naïve T cells are encountering their specific antigen presented by APCs. In Eq. 2, the numerator and the first term in the denominator indicate that higher naïve T cell frequency or higher affinity (lower *K*_i_) relative to antigen availability increases the activation rate of the clone, and the second term in the denominator accounts for interclonal competition.

Once activated, T cells expand deterministically according to a “division destiny” model (*13*), in which the number of divisions is pre-determined by the strength of the initial antigen encounter (*4*). The effector population of clone *i* at time *t* is given by:

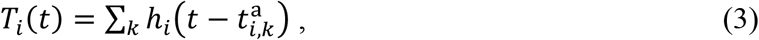

where 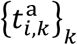are the individual activation times sampled stochastically with rate *r*_*i*_ (Eq. 2), and *h*_*i*_(*t*) describes the deterministic growth and contraction of each activated subclone:

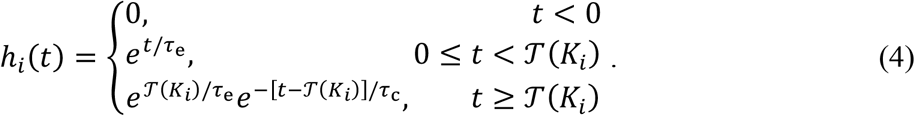

Here, consistent with Eq. 1, *τ*_e_ and *τ*_c_ are, respectively, the timescale of clonal expansion and contraction (inverse of exponential rate), and 𝒯(*K*_*i*_) is an affinity-dependent duration of expansion (e.g. higher-affinity clones expand for longer). For simplicity, we assumed that the duration of clonal scales linearly with affinity, i.e. 𝒯(*K*_*i*_) = 𝒯_0_*K*_0_/*K*_*i*_, where 𝒯_0_ is a typical duration of proliferation for T cells of reference affinity *K*_0_.

We model the viral load *V*(*t*) as undergoing logistic growth in the absence of an immune response, with elimination driven by the total population of effector T cells. The viral dynamics are thus described by

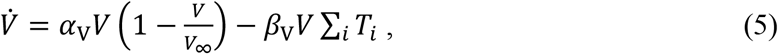

where *α*_V_ is the intrinsic replication rate of the virus, *β*_V_ is the rate of T cell-mediated clearance, and *V*_∞_ is the viral carrying capacity. This coupling allows feedback between viral growth and immune response: viral levels drive antigen presentation, which in turn activates T cells, and the expanding T cells reduce viral load. Model assumptions, parameter values, and further derivations are provided in the Supplemental Information.

### Dependence of peak abundance and peak day on precursor frequency and affinity

We used our model to ask whether variability in precursor frequency or TCR affinity could explain the lack of correlation between peak day and peak abundance observed in the experimental data (Fig. 2a,b). To this end, we simulated clonal activation within a single lymph node (LN), where approximately 100–1000 unique antigen-specific T cell clones, each with ∼10 naïve T cells, are expected to reside at any given moment (see Supplemental Information).

We first considered two cases: one in which clones varied in precursor frequency (20% variation in log-counts) while all shared the same affinity, and another in which clones had fixed precursor frequency but 20% variation in affinity. In both scenarios, we observed typical individual clonal expansion and contraction dynamics accompanied by viral clearance (Fig. 3b,c, left panels). However, the outcomes differed markedly in terms of the relationship between peak day and peak abundance. In the case of varying precursor frequency, clones peaked at different times but showed no correlation between peak time and peak abundance—consistent with our experimental findings (Fig. 3b, right panel). In contrast, variation in TCR affinity led to a strong positive correlation, with clones peaking later also reaching higher abundances due to longer proliferation periods and greater proliferative potential (Fig. 3c, right panel). When both precursor frequency and affinity were allowed to vary simultaneously (Fig. 3d), the correlation between peak day and peak abundance persisted, albeit to a slightly lesser extent.

These results argue against affinity being the primary driver of the observed variation in the timing and magnitude of T cell clonal expansion during primary infection. Specifically, the absence of correlation between peak abundance and peak day in both the experimental data and the simulations with variable precursor frequency, compared to the strong correlation observed in simulations with variable affinity, suggests that affinity alone is unlikely to account for the heterogeneity in clonal dynamics.

Differences in antigen specificity, different anatomical sites of viral replication and hence antigen presentation, and lineage identity (functional differences between CD4^+^ and CD8^+^ T cells), could also impact the observed T cell clonal dynamics. Given that infection induces a coordinated immune response across tissues and cell types, we next analyzed the data to examine whether these additional factors contribute to the timing of T cell activation and expansion.

### Matching to single-cell sequencing data reveals different kinetics between CD4^+^ and CD8^+^ T cells

To investigate whether T cell subset identity influences clonal kinetics, we cross-referenced the longitudinal clonal dynamics inferred from bulk TCR α- and β-chain sequences with matched single-cell RNA-sequencing data (*12*). Specifically, we identified CD4^+^ and CD8^+^ T cell subsets by matching exact nucleotide sequences of α- or β-chains to single-cell profiles. This yielded 27 CD4^+^ and 45 CD8^+^ clones from α-chain data, and 26 CD4^+^ and 59 CD8^+^ clones from β-chain data, across three individuals (patient numbers 4, 8, and 9).

We then compared the kinetic parameters previously inferred by fitting each clonal trajectory to Eq. 1. As shown in Fig. 4a, the clonal expansion timescale *τ*_e_ did not significantly differ between CD4^+^ and CD8^+^ T cells, while CD4^+^ clones exhibited a faster contraction timescale *τ*_c_ compared to CD8^+^ clones as assessed by β-chains (Fig. 4b). However most notably, the peak time of CD4^+^ T cells occurred substantially earlier than that of CD8^+^ T cells, a result consistently observed in both α- and β-chain matched data (Fig. 4c). In contrast, there was no significant difference in peak clonal abundance between the two subsets (Fig. 4d). These findings indicate that CD4^+^ and CD8^+^ T cells follow distinct temporal programs during primary SARS-CoV-2 infection, with CD4^+^ T cells peaking earlier and on average contracting more rapidly. These temporal differences likely mirror the distinct functional roles of CD4^+^ and CD8^+^ T cells across phases of the adaptive immune response. Motivated by these findings, we next asked whether functional differences between T cell clones could shape the kinetics of their response.

**Figure 4.**
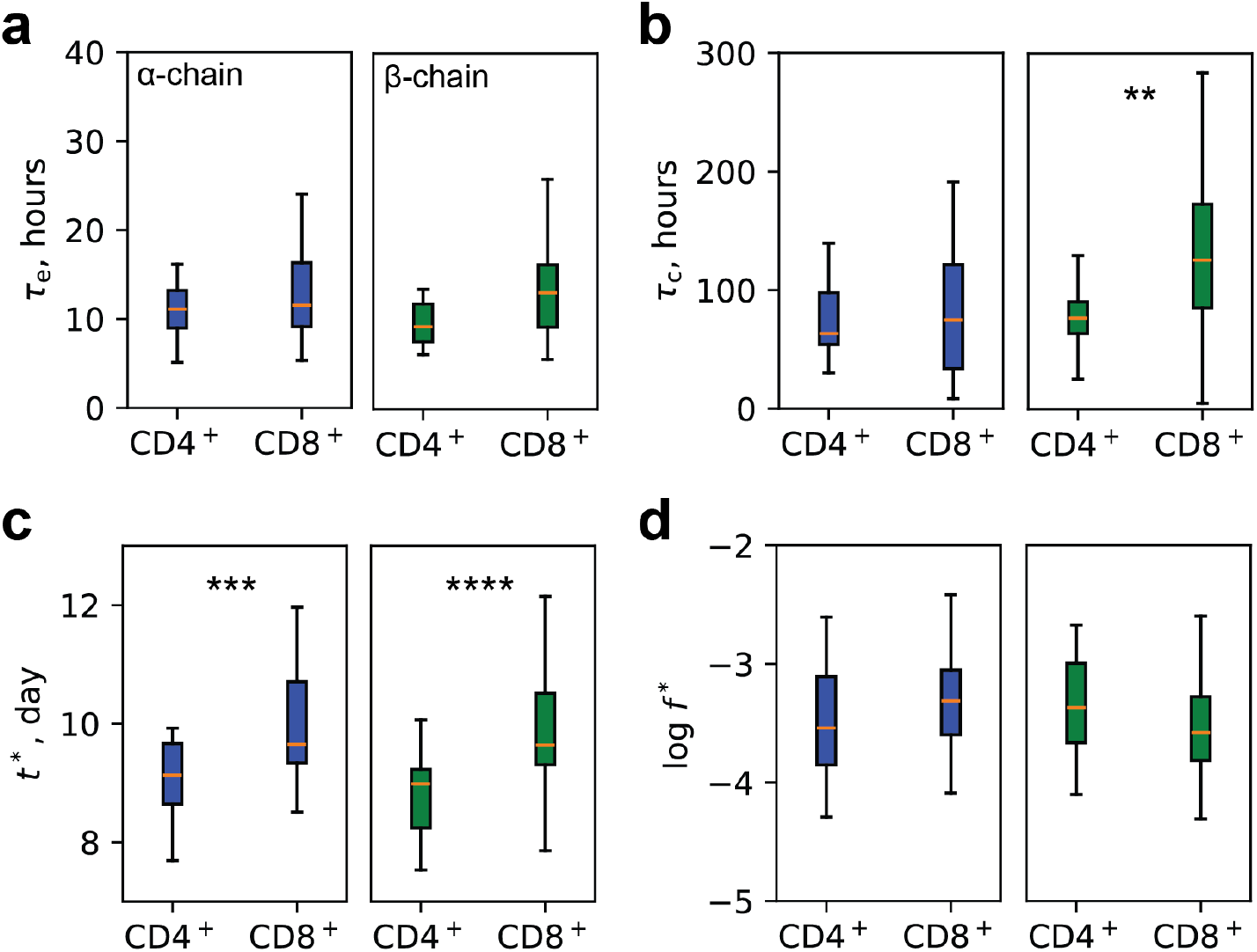
Comparison of clonal dynamics of CD4^+^ and CD8^+^ T cells. Clonal dynamics cross-matched with single-cell sequencing identified *N* = 27 CD4^+^ and *N* = 45 CD8^+^ clones by CDR3α, and *N* = 26 CD4^+^ and *N* = 59 CD8^+^ clones by CDR3β across 3 individuals. Shown are fitted parameters from Eq. 1: expansion timescale **(a)**, contraction timescale **(b)**, peak day **(c)**, and peak abundance **(d)** of expanding clones. Statistical significance was assessed with Welch’s *t*-test.

### Specificity influences the peak time of T cell clones

In particular, we hypothesized that T cells recognizing the same or similar epitopes might exhibit synchronized activation dynamics, potentially reflecting the different dynamics of different virally derived T cell epitopes. Since TCR sequence similarity often reflects shared antigen specificity, we reasoned that T cell clones with similar receptors might be activated at similar times during infection.

To test this idea, we examined pairs of expanding T cell clones within each individual and measured the difference in their peak days, defined as 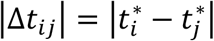for clone labels *i* and *j*. In order to estimate common specificity, we classified a pair of clones as “matched” if they shared the same V gene and J gene, and as “unmatched” otherwise. While this definition does not guarantee shared specificity, it provides a workable proxy. This analysis was performed separately for the α-chain and β-chain sequences, and restricted to individuals for whom we could identify at least 10 unique matched pairs of expanding clones. Three individuals met this criterion for both α-chain and β-chain analyses.

We found that in both α- and β-chain analysis, 2 out of 3 individuals showed a statistically significant reduction in |Δ*t*^∗^| for matched pairs, while the remaining one also showed a reduction that did not reach statistical significance (Fig. 5a). These results suggest that T cells with similar TCR sequences—likely recognizing overlapping epitope—tend to expand with more synchronized kinetics. While one alternative explanation is that the reduction in |Δ*t*^∗^| arises from systematic differences between CD4^+^ and CD8^+^ T cell expansion dynamics, the coincidence rate of V gene matches across lineages was only slightly elevated, making this unlikely (Fig. S3). This prompted us to consider whether the observed coordination instead reflects variation in viral antigen availability across tissue sites.

**Figure 5.**
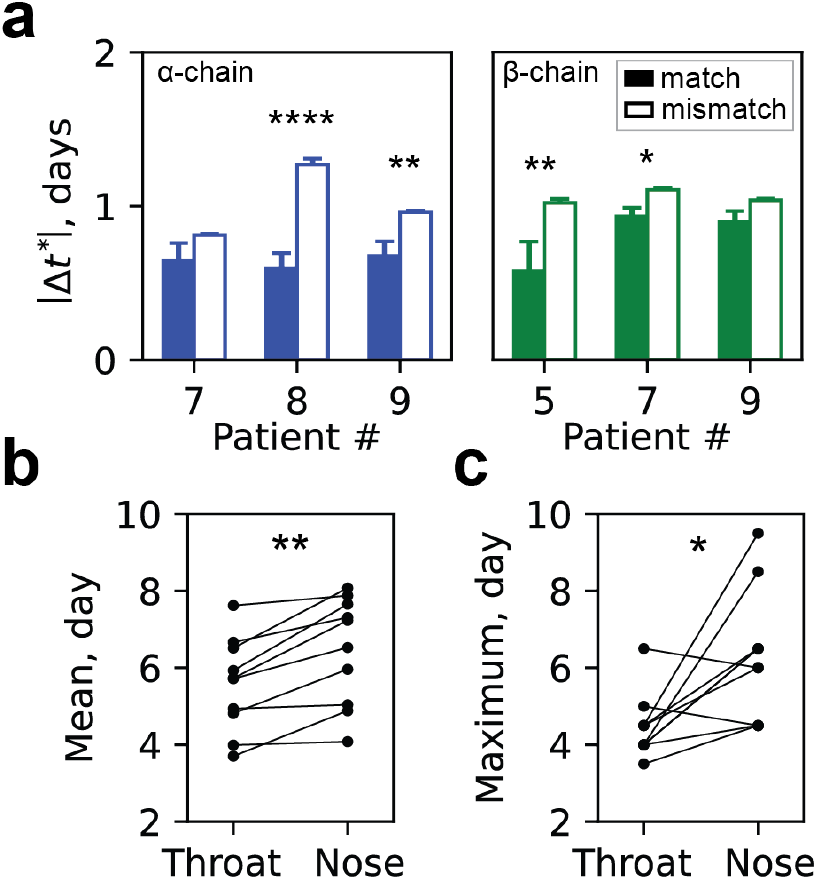
Synchronized kinetics of expanded T cell clones. **a**, Difference in peak day between pairs of expanding T cell clones, stratified by whether they share the same V and J genes (filled bars) or not (empty bars). Only individuals with at least five clone pairs sharing both V and J genes are included. Expanding T cell clones identified from α-chain sequences are shown in blue (left panel), while those from β-chain sequences are shown in green (right panel). Statistical significance was assessed using Welch’s *t*-test. **b**,**c**, Viral loads in the throat and nose from the same 10 patients, summarized by the mean **(b)** and maximum **(c)** of their viral load distributions over 28 days post inoculation. Statistical significance was determined by paired t-test.

### Viral load kinetics vary across anatomical sites

Because T cell activation is initiated by antigen presentation, which in turn depends on local viral dynamics, we asked whether variation in viral kinetics across tissue sites could contribute to the heterogeneity in T cell activation timing. To this end, we analyzed the kinetics of viral load in both the throat and nose of all 10 individuals, with measurements taken every 12 hours over the first 20 days following SARS-CoV-2 inoculation.

We characterized viral kinetics using two measures: the first moment (mean) of the viral load distribution and the time of peak viral load. As shown in Fig. 5b,c, and consistent with previous analysis, both the average first day of viral detection and the day of maximal viral load occurred earlier in the throat than in the nose (*11*). Given that the nasal cavity and nasopharynx drain to distinct LNs (*14, 15*), this observation suggests spatially staggered immune activation, with different LNs becoming engaged at different times during infection. This spatiotemporal variation in viral kinetics prompted us to ask how antigen-specific T cell responses evolve over time, particularly with respect to memory formation.

### Persistent T cells across antigen-specific clones and individuals

To investigate memory formation, we used the persistence of antigen-specific T cell clones measured by their abundance at later timepoints as a proxy. In prior sections, we characterized the kinetics of T cell clones using fits to Eq. 1, which focuses on the behavior around the peak. However, we noticed that some antigen-specific clones deviated from this fit at later timepoints, suggesting a secondary timescale of decay (Fig. S2). This slower phase is likely associated with entry into the memory compartment, which has been characterized by a shift in decay kinetics after initial contraction (*10, 16*).

To assess memory-like persistence, we examined the abundance of expanding clones at day 28 post-inoculation. On average, 50-60% of expanding clones remained detectable at this timepoint, though this fraction varied widely across individuals, ranging between 25–75% (Fig. 6a). To further characterize persistence, we defined two metrics. First, we computed the log fold-change in abundance at day 28 relative to the peak, log *f*_D28_/*f* ^*^, which quantifies the degree of clonal contraction (Fig. 6b). Across individuals, we observed an approximately 10-fold decrease in abundance from peak to day 28 (Fig. 6c). However, the extent of persistence could not be predicted from earlier clonal kinetics; we found no significant correlation between any of the kinetic parameters (e.g., peak abundance, expansion rate, contraction rate, or peak timing) and the level of persistence at day 28 (Fig. S4).

**Figure 6.**
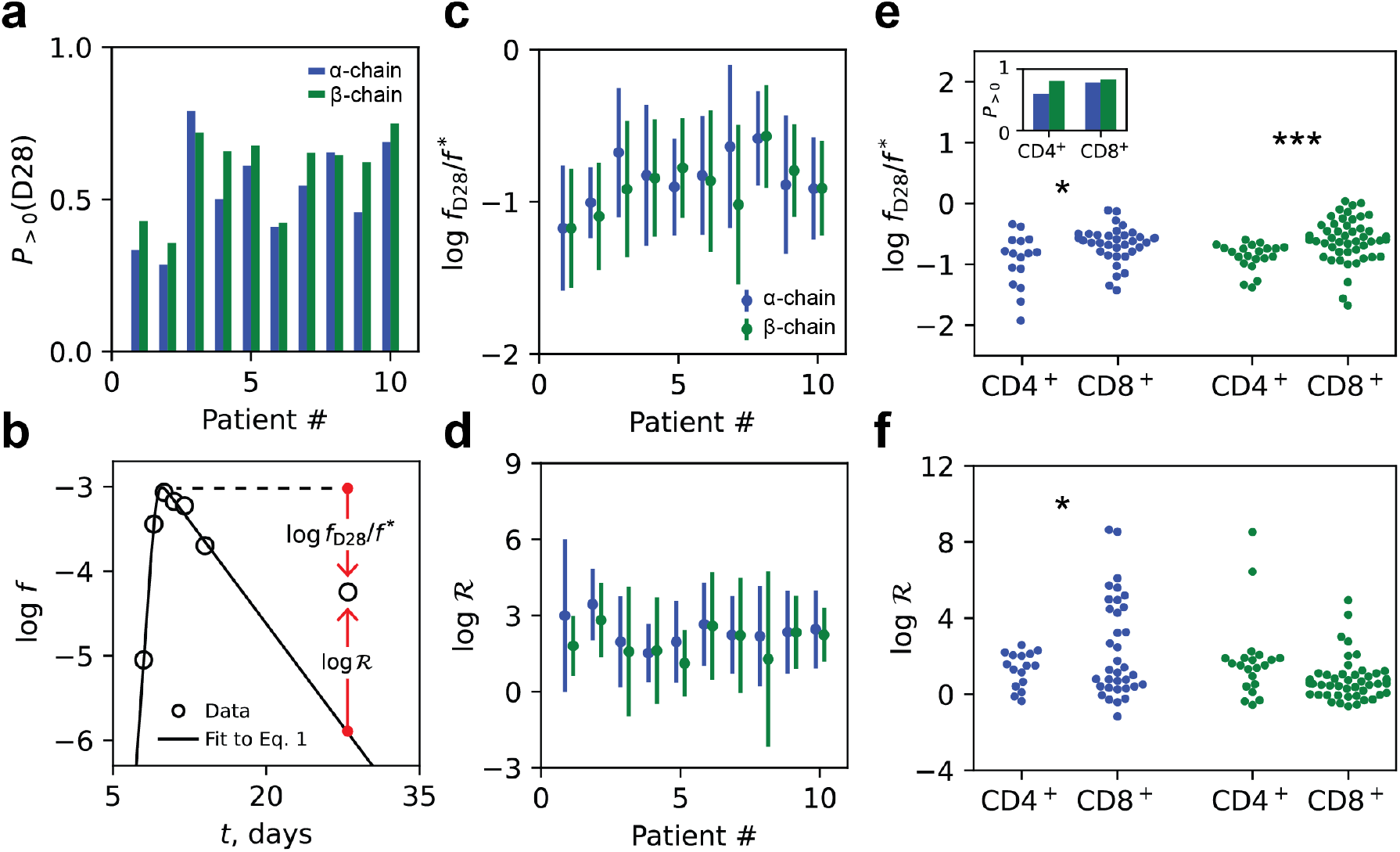
Persistence of expanding clones varies across individuals and lineages. **a**, Fraction of expanding clones remaining at detectable levels on day 28 after inoculation, shown for each of the 10 individuals. **b**, Schematic illustration of T cell persistence metrics. Persistence of individual clones was quantified using two additional metrics: the log fold-change in abundance from peak to day 28, log *f*_D28_/*f* ^*^, and the deviation from model-predicted abundance at day 28 based on fits to Eq. 1, denoted log *ℛ*. **c**,**d**, Distribution of persistence metrics across individuals. Shown are log *f*_D28_/*f* ^*^ **(c)** and log *ℛ* **(d)**, with error bars indicating standard deviation. **e**,**f**, Comparison of persistence metrics between CD4^+^ and CD8^+^ T cells across three individuals with matched single-cell data. log *f*_D28_/*f* ^*^ **(e)** and log *ℛ* **(f)** are shown. Inset in **(e)** shows the fraction of persistent clones by lineage. Statistical significance was assessed using Welch’s *t*-test.

Second, we analyzed the deviation, which we denote as log *ℛ*, between observed and model-predicted abundance at day 28, based on extrapolation from fits to Eq. 1 (Fig. 6b). This measure revealed greater variability both within and across individuals, spanning several orders of magnitude (Fig. 6d). Assuming that T cell clones persisting at detectable levels to day 28 primarily reflect the memory compartment, these results suggest that the rate of conversion to memory varies substantially across clones and individuals, highlighting the heterogeneous and likely context-dependent nature of memory formation during primary infection. Building on the observed variability in clonal persistence across individuals, we wondered whether these differences could be explained by T cell lineage.

### Persistent CD4^+^ and CD8^+^ T cells exhibit distinct long-term contraction patterns

To determine whether T cell lineage influences clonal persistence, we analyzed the persistence metrics defined in the previous section for cross-referenced CD4^+^ and CD8^+^ T cell clones across the three individuals with matched single-cell data. The overall fraction of expanded clones with detectable abundance at day 28 did not differ significantly between CD4^+^ and CD8^+^ T cell subsets (Fig. 6e, inset). Among the persistent clones, CD4^+^ T cells exhibited significantly lower abundance relative to their peak, as measured by log *f*_D28_*/f* ^*^ (Fig. 6e), but an increased median abundance relative to the abundance predicted from the post-peak contraction kinetics (Fig 6f). Taken together, the data are consistent with a faster initial contraction of CD4^+^ T cells, with a lower maintenance frequency of memory cells.

## Discussion

The SARS-Cov-2 challenge study carried out in virus-naïve individuals in 2020 provided an opportunity to study the dynamics of a human primary immune response under conditions of exceptional synchronicity and temporal sampling density. We quantified the clonal dynamics of antigen-specific T cells during primary SARS-CoV-2 infection and found remarkable consistency at the population level across 10 individuals, despite natural variation in their TCR repertoires. By longitudinally tracking clonal abundance, we captured key features of the T cell response— specifically, the timescales of expansion and contraction, and the timing and magnitude of each clone’s peak. Our analysis shows that T cell clones expand at a near-maximal proliferation rate, consistent with previous estimates. In contrast, contraction rates were slower and more variable across clones, suggesting heterogeneous clonal fates.

Due to limits in sequencing depth, our data primarily capture clonal trajectories around their peak when the clonal abundance is above a threshold level. To infer possible upstream drivers of clonal activation and expansion, we employed a mathematical model to test the predicted effects of variation in T cell precursor frequency and the strength of antigenic stimulus, itself a combination of TCR affinity and the concentration of presented antigen. We found that, in the model, variation in precursor frequency, rather than affinity alone, predicted the experimentally observed lack of correlation between peak day and peak abundance. This suggests that precursor frequency could play an important role in specifying the early clonal dynamics of individual T cell clones in primary infection. Significant variation in precursor frequency within the naïve T cell population has been observed previously in a general context (*17*) and proposed to provide differential early responses in SARS-Cov-2 and LCMV infection (*18*).

Interestingly, this result may reflect a system-level principle: maintaining consistent clonal peak amplitudes over a period of several days may be beneficial for robustness, ensuring broad coverage of diverse epitopes without excessive amplification of any single clone. Nonetheless, our findings do not rule out more complex, unmodeled sources of variation—such as differences in antigen availability, tissue microenvironment, or heterogeneity in activation conditions including co-stimulatory signals and cytokine profiles—leaving important avenues for future investigation.

We then investigated whether clonal kinetics differ by T cell lineage. By integrating single-cell transcriptomic data, we found that CD4^+^ T cells peaked earlier and contracted more rapidly than CD8^+^ T cells. This is consistent with their distinct roles in viral infection: CD4^+^ T cells act earlier to coordinate the immune response and provide help to other immune cells, while CD8^+^ T cells persist to ensure cytotoxic clearance. It is difficult to compare these estimates reliably with previous studies because, at least in the human, data with the temporal precision and resolution to capture these very transient rapid peaks is rarely available. Thus, although previous studies in mice have reported more extensive clonal expansion in CD8^+^ T cells compared to CD4^+^ T cells (*10, 19, 20*), we did not observe a consistent bias in maximum expansion magnitude between these subsets in our human viral infection cohort (Fig. 4d). Further studies will be needed to determine the generality of our results, and whether the relative expansion dynamics are shaped by pathogen-specific or other context-dependent factors.

To probe additional factors which may shape T cell clonal dynamics, we analyzed TCR sequence similarity and found that clones with similar sequences tend to peak at similar times, suggesting coordinated dynamics between T cell clones of similar inferred specificity. We also noted that the peak in viral titers in the two sampled anatomical site, the nose and throat epithelium, are not synchronous but tend to peak first in the throat. Therefore, synchronous expansion of co-specific TCRs may result from staggered LN engagement as naïve T cells with similar specificity could preferentially accumulate in different draining LNs perhaps due to selective local self-antigen presentation (*21*), or might encounter distinct antigens at varying times due to localized presentation by diverse APCs (*22*). These spatiotemporal differences in antigen exposure and co-stimulatory context could result in the structured activation timing observed for T cell clones with similar sequences. Overall, these results support the idea that spatial organization of the immune response—mediated through lymphatic architecture and antigen spread—imposes structure on T cell clonal dynamics.

Observed difference in the contraction phase may reflect underlying differences in memory cell differentiation. Using clonal persistence at day 28 as a proxy for memory formation, we found that approximately 55% of antigen-specific clones remained detectable at this timepoint, though this fraction varied widely across individuals (25–75%). Among these persistent clones, we observed an average ∼10-fold reduction in abundance relative to the peak, accompanied by broad deviations from early contraction kinetics (Fig. 6e,f). Strikingly, the degree of persistence was not significantly correlated with early kinetic parameters—such as expansion rate, peak timing, or peak abundance (Fig. S4)—consistent with memory conversion being governed by intrinsic differentiation programs rather than competitive dynamics during the expansion or early contraction phase. Although there was no significant difference between CD4^+^ and CD8^+^ T cells in the proportion of responding clones which were still detectable at day 28, the frequency of CD4^+^ cells was smaller, perhaps reflecting a lower maintenance frequency of CD4^+^ cells memory cells compared to CD8^+^ cells.

Together, these results highlight how both cell-intrinsic and population-level factors, such as TCR specificity, lineage identity and precursor frequency, and extrinsic spatiotemporal features including viral dynamics, LN engagement and tissue-specific antigen presentation, contribute to the structure and variability of the T cell response during primary infection. Our findings underscore the importance of considering not only the intrinsic properties of individual T cells, but also the broader anatomical and temporal context in which they operate, as well as their lineage identity. This integrated perspective could guide the development of next-generation vaccines aimed at more precisely recruiting and sustaining specific memory T cell lineages.

## Materials and Methods

### SARS-CoV-2 controlled human infection model

The details of the study have been published previously (*11*). Briefly 36 healthy adults aged 18 to 30 years were recruited to this single-center, phase 1, open-label, first-in-human study. Participants had no previous SARS-CoV-2 vaccination or documented infection and were excluded if positive for SARS-CoV-2 anti-S antibodies measured at screening by MosaiQ COVID-19 antibody microarray (Quotient). Two participants were excluded because of seroconversion between screening and inoculation resulting in 34 seronegative individuals for analysis. The study was conducted in accordance with the protocol; the consensus ethical principles derived from international guidelines, including the Declaration of Helsinki and Council for International Organizations of Medical Sciences International Ethical Guidelines; applicable ICH Good Clinical Practice guidelines; and applicable laws and regulations. The screening protocol and main study were approved by the U.K. Health Research Authority’s Ad Hoc Specialist Ethics Committee (references 20/UK/2001 and 20/UK/0002) and registered with ClinicalTrials.gov (identifier NCT04865237). The study was conducted in a high-containment clinical trials unit at the Royal Free London NHS Foundation Trust where participants were housed in single-occupancy, negative pressure side rooms.

Participants were inoculated intranasally by pipette with 10 TCID50 of a Good Manufacturing Practice wild-type SARS-CoV-2 strain (20A clade of the B.1 lineage having the D614G mutation; GenBank accession number OM294022) divided between both nostrils (100 μl each). Safety was assessed with daily blood tests, spirometry, electrocardiograms, clinical assessments (vital signs, symptom diaries, and clinical examination), and a CT scan of the chest on day 5 (all participants) and day 10 (infected participants only).

### Sample collection

3 ml whole blood for T cell receptor (TCR) repertoire sequencing (2.5 ml) was collected daily from each volunteer for the first 14 days and then at day 28 in RNA-stabilizing tubes (PAXgene).

Virological assessments of infections were based on 12-hourly mid-turbinate and throat flocked swabs placed in 3 ml of the viral transport medium (BSV-VTM-001, Bio-Serv) that was aliquoted and stored at −80°C. Aliquots were analyzed by RT-PCR and quantitative culture by FFA as previously described.

### TCR sequencing

17 individuals showed sustained viremia after inoculation, which returned to baseline in all individuals by day 28. TCRseq was carried out at days -1, 1, 4, 7, 8, 9, 10, 11, 12, 14, and 28 for all individuals who showed evidence of sustained infection (at least 3 PCR+ adjacent viral measurements). The α- and β-chains of the TCR were sequenced using a quantitative experimental and computational pipeline described previously (*23*–*25*). The pipeline introduces unique molecular identifiers attached to individual cDNA molecules which allows correction for sequencing error PCR bias, and provides a quantitative and reproducible method of library preparation. The Decombinator software used to annotate TCRs and perform error correction is freely available at https://github.com/innate2adaptive/Decombinator.

### Constructing longitudinal trajectories of clonal abundance

Each sequenced blood sample yielded hundreds of thousands of unique TCR sequences for both the α- and β-chains. Longitudinal clonal trajectories were constructed by identifying TCRs with identical CDR3 junction nucleotide sequences and matching V and J gene usage across different timepoints. To enable comparison across samples and timepoints with varying sequencing depths, counts were normalized by the total number of TCR reads in the respective sample, yielding a measure of relative abundance.

### Identification of expanding T cell clones

A T cell clone was classified as expanding if its longitudinal abundance satisfied the following criteria: (i) the clone was not detected on days –1 and 1; (ii) it exhibited a statistically significant increase in abundance, assessed using a one-sided exact Poisson rate comparison, on at least two consecutive days between days 7 and 12; and (iii) when fit to Eq. 1, the inferred peak day of abundance fell between days 7 and 14.

To determine whether a clone exhibited a significant increase in abundance across two consecutive time points, we assumed that the number of reads for a given clone on day *i* follows a Poisson distribution: 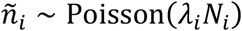, where λ_*i*_ is the true underlying frequency of the clone on day *i*, and *N*_*i*_ is the total number of T cells observed in the sample from that day.

Under the null hypothesis that the clone’s abundance remains unchanged between two days (i.e., *H*_0_: λ_*i*_ = λ_j_ = λ), the total count over the two days, 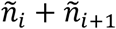, is distributed as: 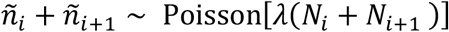. Conditioning on the total count, the number of reads on day *i+*1 follows a binomial distribution:

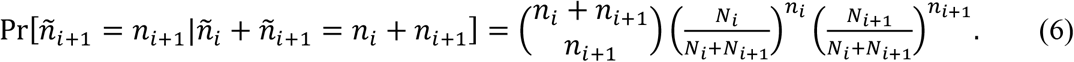

This provides the exact conditional distribution under the null. The corresponding one-sided *p*-value is given by:

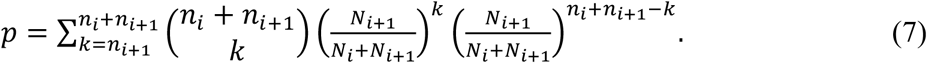

We rejected the null hypothesis and considered the increase to be statistically significant if *p* < 0.05.

### Cross-referencing bulk and single-cell sequencing data

The single-cell data has been published previously (*12*). Expanded clones identified from bulk TCR sequencing were cross-matched to single-cell data by comparing the CDR3 junction amino-acid sequence and V gene identity within the same individual. In cases where multiple single-cell TCRs matched a given bulk TCR clone (i.e., shared the same CDR3 junction amino-acid sequence and V gene), we examined the lineages (e.g., CD4^+^ vs. CD8^+^) assigned to each of the matched single-cell clones. The majority lineage— the lineage represented by the largest number of matching single-cell clones—was assigned to the bulk clone. In instances where there was a tie between lineages (i.e., equal number of CD4^+^ and CD8^+^ matches), the clone was excluded from lineage assignment to avoid ambiguity.

## Acknowledgements

The authors gratefully acknowledge support from the UK Vaccine Taskforce of the Department of Business, Energy and Industrial Strategy of Her Majesty’s Government (BEIS); the Wellcome Trust (grant ref: 224530/Z/21/Z); the Rosetrees Foundation; and the NIHR UCL Biomedical Research Centre (BRC). This study was supported by the Natural Sciences and Engineering Research Council of Canada and Stanford Science Fellowship (to A.G.T.P.); the Chan Zuckerberg Initiative DAF (grant number DAF2024-342781), an advised fund of the Silicon Valley Community Foundation (to N.S.W. and A.G.T.P.); Princeton University through the Center for the Physics of Biological Function (to N.S.W. and A.G.T.P.); Princeton University Visiting Fellowship (to B.C.); and the NIHR Imperial BRC award to Imperial College Healthcare NHS Trust and Imperial College London (to C.C.). A.G.T.P thanks Julia Merkenschlager for helpful discussions and critical reading of the manuscript.

We also thank the following organizations for their invaluable contributions to the development and implementation of the SARS-CoV-2 human challenge project: the Royal Free London NHS Foundation Trust; the Human Infection Challenge Network for Vaccine Development (HIC-Vac); NIHR Clinical Research Network staff at the Royal Bolton Hospital; and ISARIC4C Investigators (https://isaric4c.net/about/authors/) for providing the clinical material for challenge virus production. ISARIC4C is funded by the National Institute for Health Research (NIHR; award CO-CIN-01), the Medical Research Council (MRC; grant MC_PC_19059), the NIHR Health Protection Research Unit in Emerging and Zoonotic Infections at the University of Liverpool in partnership with Public Health England (PHE), in collaboration with the Liverpool School of Tropical Medicine and the University of Oxford (NIHR award 200907). The Liverpool Experimental Cancer Medicine Centre provided infrastructure support for this research (grant reference C18616/A25153), and the NIHR Health Protection Research Unit in Respiratory Infections also provided support (NIHR award 200927).

The views expressed are those of the authors and not necessarily those of the NHS, the NIHR, the DHSC, or BEIS.

## Author contributions

Conceptualization, A.G.T.P., N.S.W., and B.C.; methodology, A.G.T.P., C.G.C., A.T.M., N.S.W., and B.C.; Investigation, A.G.T.P., N.S.W., and B.C.; writing—original draft, A.G.T.P., N.S.W., and B.C.; writing—review & editing, All authors; funding acquisition, N.S.W., and B.C.; resources, A.C., J.R., C.T., L.B., G.N., S.B., B.K., M.K., A.B., A.C., R.G.H.L., M.Z.N., S.A.T., H.W., M.N., and C.C.; supervision, N.S.W and B.C.

## Data and code availability

This paper analyzes existing, publicly available data, accessible at https://doi.org/10.1038/s41591-022-01780-9, and https://doi.org/10.1038/s41586-024-07575-x.

All original code has been deposited at Zenodo at [DOI] and is publicly available as of the date of publication.

Any additional information required to reanalyze the data reported in this paper is available from the lead contact upon request.

## Supplemental information

Document S1. Figures S1–S4, Table S1, and supplemental references

## Supplemental Information

### 1. Mathematical model

In this section, we outline the model, which closely follows the framework developed by de Boer and Perelson [1].

We consider a scheme where, in a given lymph node, a naive T cell of clonotype *i*, denoted 𝒩_*i*_, interacts with a peptide-loaded binding site on antigen-presenting cells (APCs), denoted 𝒮, to form an interaction complex 𝒞_*i*_, with association and dissociation rates 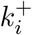 and 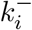, respectively. Upon complex formation, naive T cells can be activated at rate 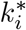, becoming an activated T cell, denoted 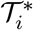. This process is summarized by:

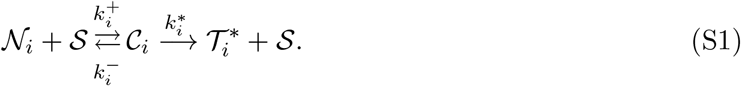

We denote the total naive T cell population in clonotype *i* (including both free T cells 𝒩_*I*_ and complexes 𝒞_*i*_) by 𝒩_*i*_. Its dynamics are:

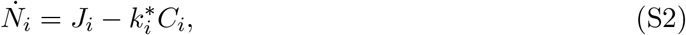

where *J*_*i*_ denotes the influx of naive T cells, and *C*_*i*_ is the number of complexes. The activated T cell population, denoted 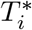, evolves as:

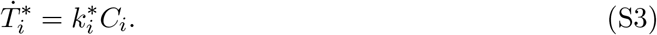

Activated T cells proliferate and differentiate into effector T cells *T*_*i*_, which contribute to viral clearance:

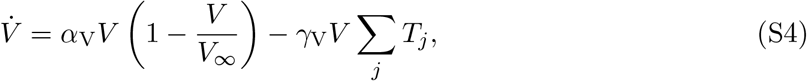

where *V* is the viral load, *α*_V_ is the viral replication rate, and *γ*_V_ is the rate of virus elimination by T cells.

To model antigen presentation, let 𝒮_free_ represent unoccupied peptide-presentation sites in a given lymph node. Presentation of viral particles 𝒱 to these free sites proceeds via

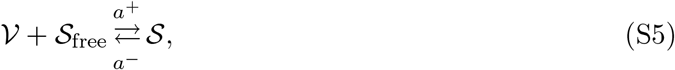

where *a*^+^ is the rate of antigen presentation, and *a*^−^ is the decay rate of presented antigen.

Within a given lymph node, let *S*_tot_ denote the total number of potential peptide-presentation sites, and define *S*_free_ = *S*_tot_ − *S* as the number of unoccupied sites. Here, *S* denotes the total number of sites currently presenting peptide, including both unbound antigen-presenting sites 𝒞 and those bound to T cells of any clonotype (i.e. Σ_*i*_ *C*_*i*_). The dynamics of antigen presentation is then given by

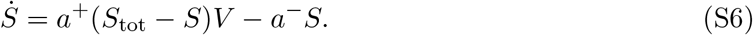

#### 1.1 Quasi-steady state approximations

##### 1.1.1 Antigen presentation

Assuming quasi-steady state 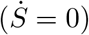, the total number of peptide-loaded sites becomes:

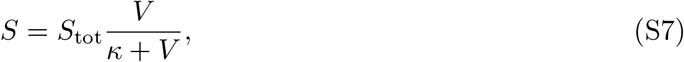

where *κ* ≡ *a*^−^*/a*^+^ represents the inverse of antigen presentation efficiency.

##### 1.1.2 Interaction complexes

The dynamics of *C*_*i*_ are:

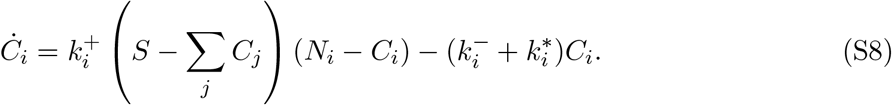

Assuming a quasi-steady state for each *C*_*i*_, and expanding to first order in *C*_*i*_, i.e. ∼ *𝒪* (*C*_*i*_), we obtain:

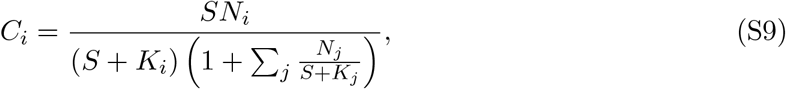

where 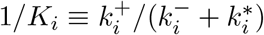 characterizes the affinity of T cells for the antigen.

##### 1.1.3 Naive T cell balance

To simplify, we assume that influx exactly balances activation 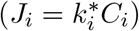, so the precursor pool remains constant. This gives:

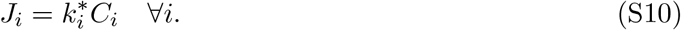

Thus, the activation rate depends on the viral load via *S*(*V*):

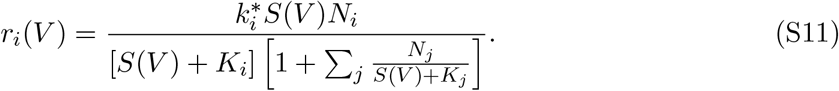

#### 1.2 Stochastic activation via inhomogeneous Poisson process

We model activation of naive T cells as a discrete stochastic process in continuous time. For each clonotype *i*, activation events occur at random times 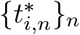, where each event corresponds to the transition of a naive T cell into an activated state. We assume that these activation times are generated by an inhomogeneous Poisson process with a time-dependent rate *r*_*i*_(*t*), which depends on antigen availability and TCR affinity.

##### 1.2.1 Definition of activation indicator

Let *r*_*i*_(*t*) denote the instantaneous activation rate for clone *i*. The probability that an activation occurs in the infinitesimal time interval [*t, t* + *dt*) is given by

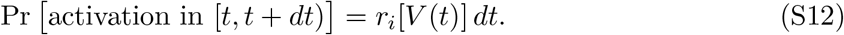

The corresponding activation indicator function is defined as

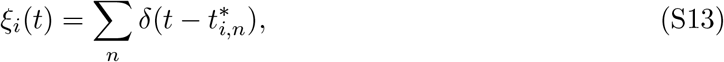

where *δ*(·) is the Dirac delta function, and 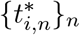 are the (random) activation times for clone *i*.

Each term 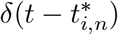 represents a discrete activation event occurring at time 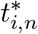 .

##### 1.2.2 Effector T cell dynamics and expectations

Following activation, a T cell undergoes deterministic proliferation governed by a response function *h*(*t, K*_*i*_) that depends on the time since activation and the affinity 1*/K*_*i*_. The effector population for clone *i* then evolves as:

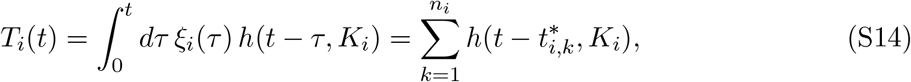

where *n*_*i*_ is the number of activation events of naive T cells of clonotype *i*. The above expression sums the contributions from all past activation events.

Taking the expectation over realizations of the stochastic process, and using the fact that 𝔼 [*ξ*_*i*_(*t*)] = *r*_*i*_(*t*), we obtain the expected rate of effector accumulation:

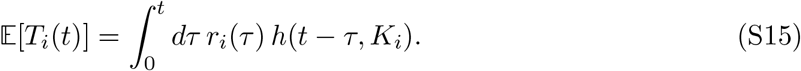

It also follows that 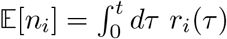.

##### 1.2.3 Deterministic proliferation

Upon activation, we assume that the activated T cell proliferates at a fixed rate, *α*_T_, for an affinity-dependent duration 𝒯 (*K*_*i*_) and then the resulting lineage decays at a fixed rate *β*_T_. The corresponding response function is

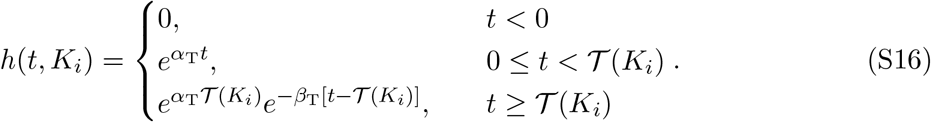

A simple choice for the duration of proliferation is

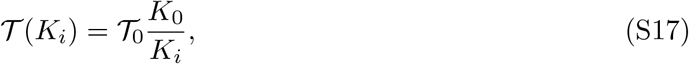

where 𝒯_0_ is a typical duration of proliferation for T cell with affinity 1*/K*_0_.

### 2 Parameters

#### 2.1 Number of naive T cells

There are ∼ 10^11^ naive T cells in humans [2], where a fraction of 10^−6^ − 10^−5^ are specific to any given antigen [3]. Given that there are ∼ 10^8^ unique clonotypes of naive T cells in humans [4, 5], this implies that for a particular antigen there are *N*_clone_ ∼ 10^2^ − 10^3^ specific T cell clones, with average ∼ 10^3^ naive T cells per clone.

The average number of T cells in the same clonotype that is present in a specific LN can be estimated as

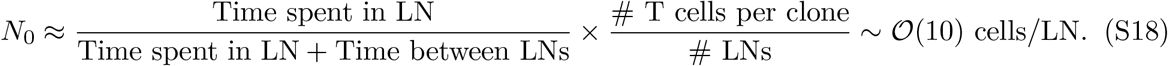

##### 2.1.1 Interaction kinetics

Assuming that at most ∼ 10% of naive T cells are interacting with a presented antigen at any given time (i.e., in the limit *V* ≫ *κ*), we use Eq. S9 to estimate the typical number of interacting cells:

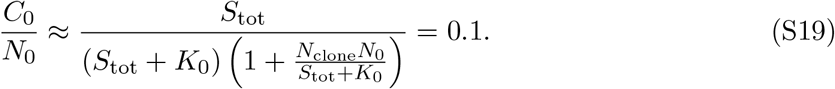

Solving for 1*/K*_0_ yields

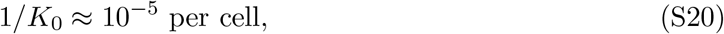

which is consistent with the estimates of de Boer and Perelson [1].

Similarly, DC-T cell interaction typically lasts 3−4 minutes [6]. Assuming productive interactions are rare, such that 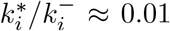 corresponding to approximately 1% of DC-T cell interactions leading to activation [1], we have that

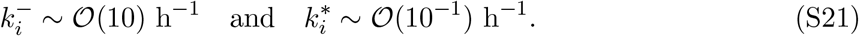

### 3 Figures and tables

**Table S1.**
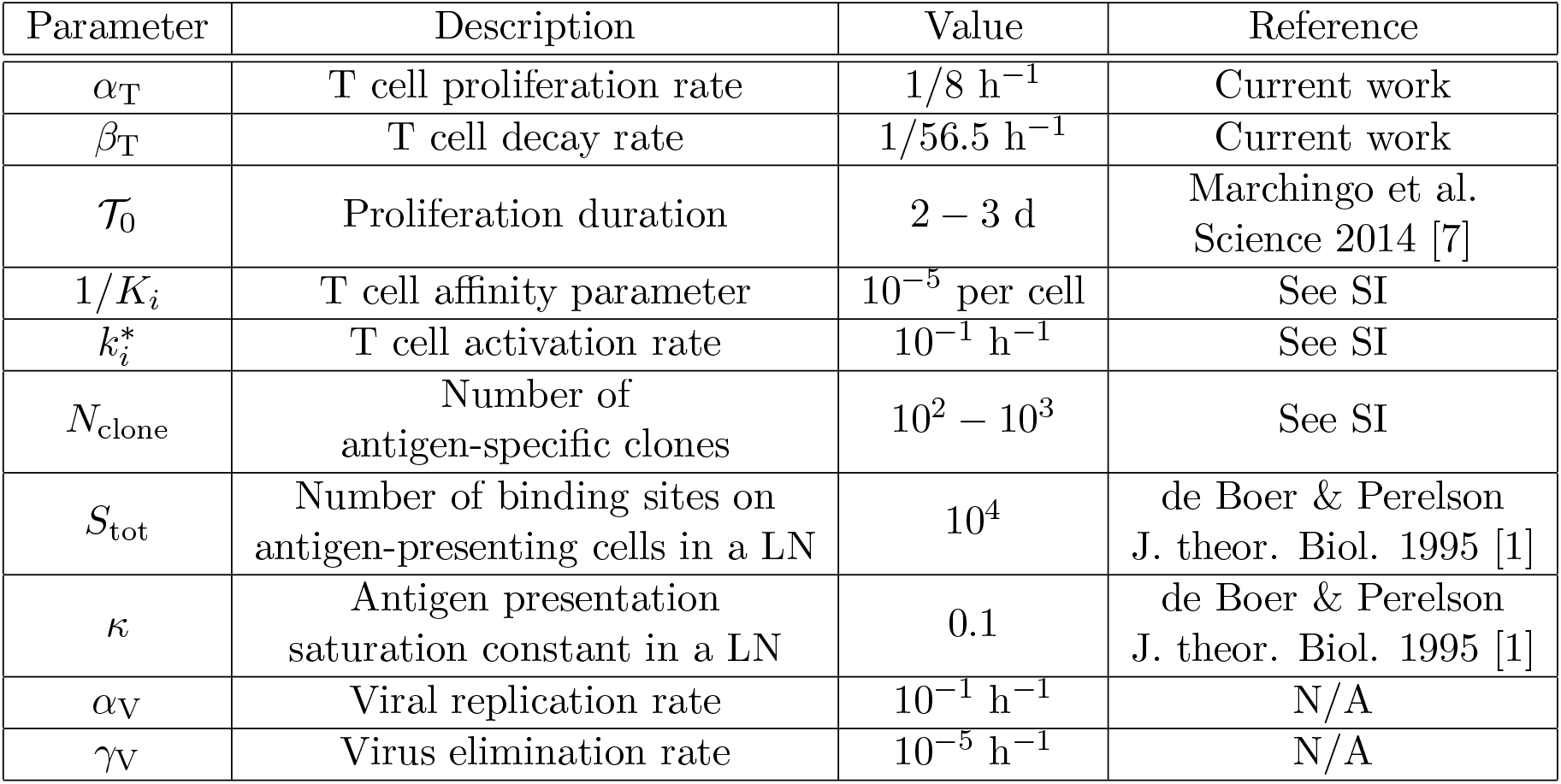
Simulation parameters.

**Figure S1.**
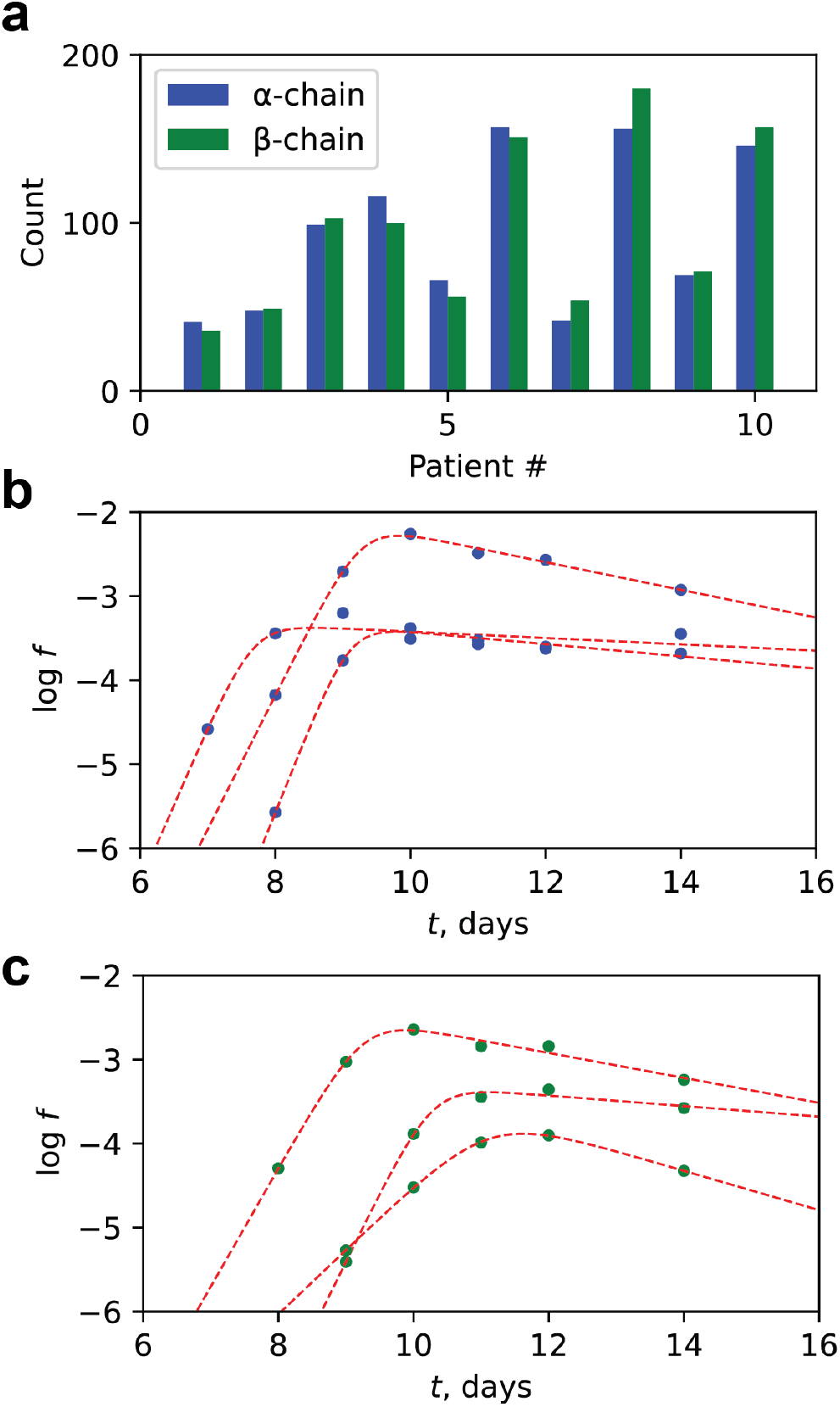
Identification and characterization of expanding T cell clones. **a**, Number of expanding clones identified in each individual. **b**,**c**, Examples of longitudinal clonal abundance data constructed between day 7 and day 14 for representative *α*-chain (**b**) and *β*-chain (**c**) TCR clones. Solid dots represent observed data points; dotted lines indicate model fits.

**Figure S2:**
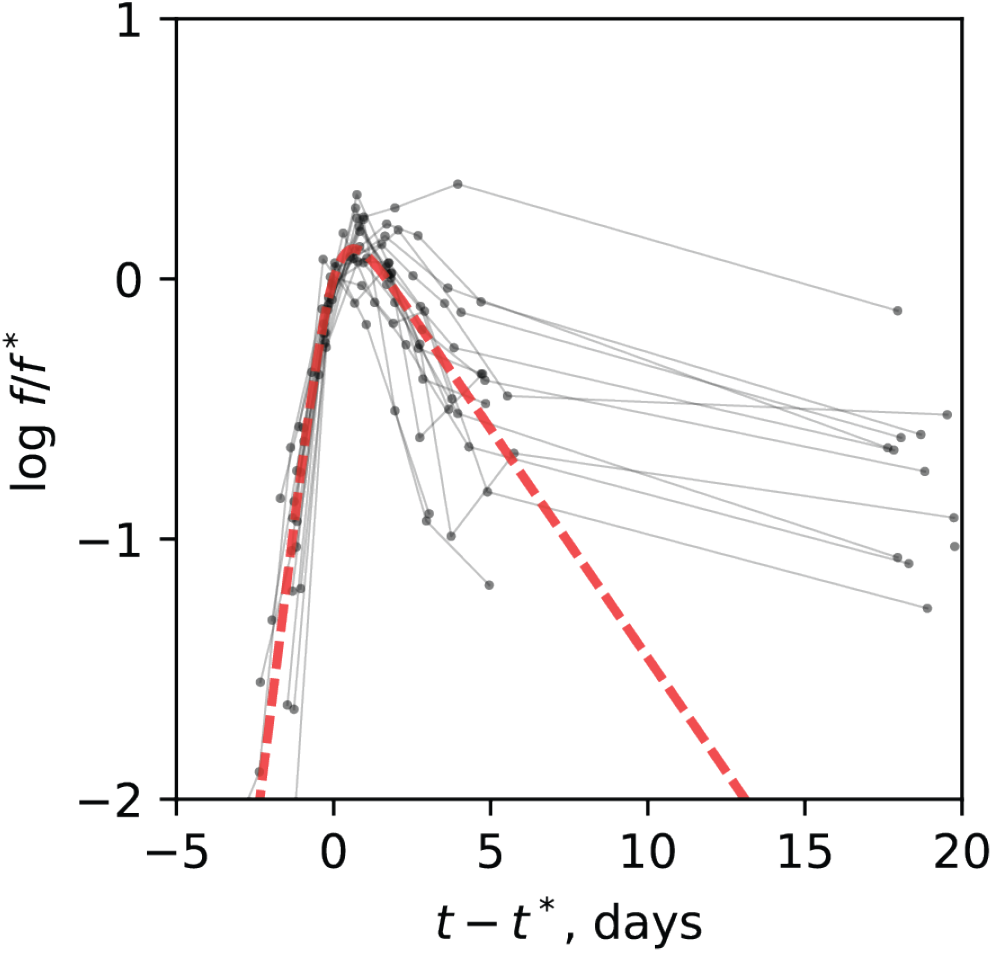
Secondary contraction timescale. Example longitudinal clonal abundance trajectories are shown in black. The red line represents the average fitted trajectory around the peak to guide the eye.

**Figure S3.**
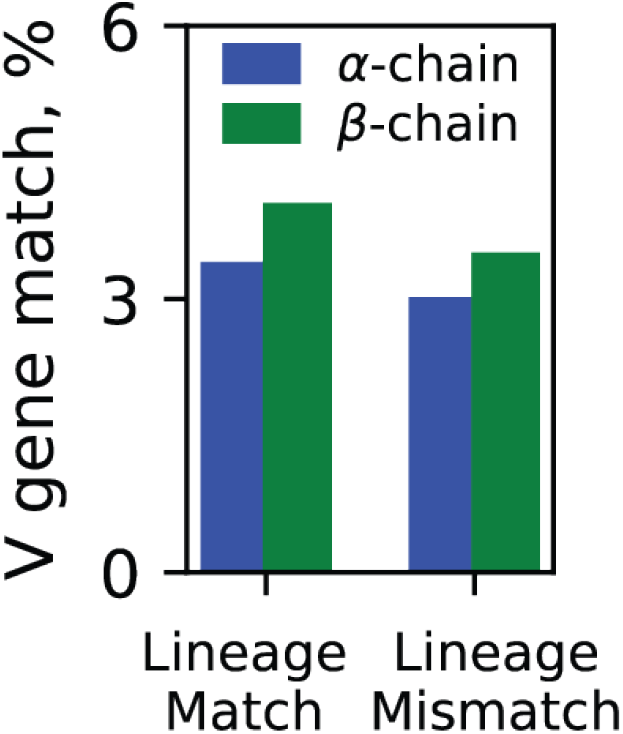
Percentage of unique TCR clone pairs within each individual that shared the same V gene, conditioned on whether the pair was lineage-matched (both CD4^+^ or both CD8^+^) or lineage-unmatched (one CD4^+^, one CD8^+^).

**Figure S4.**
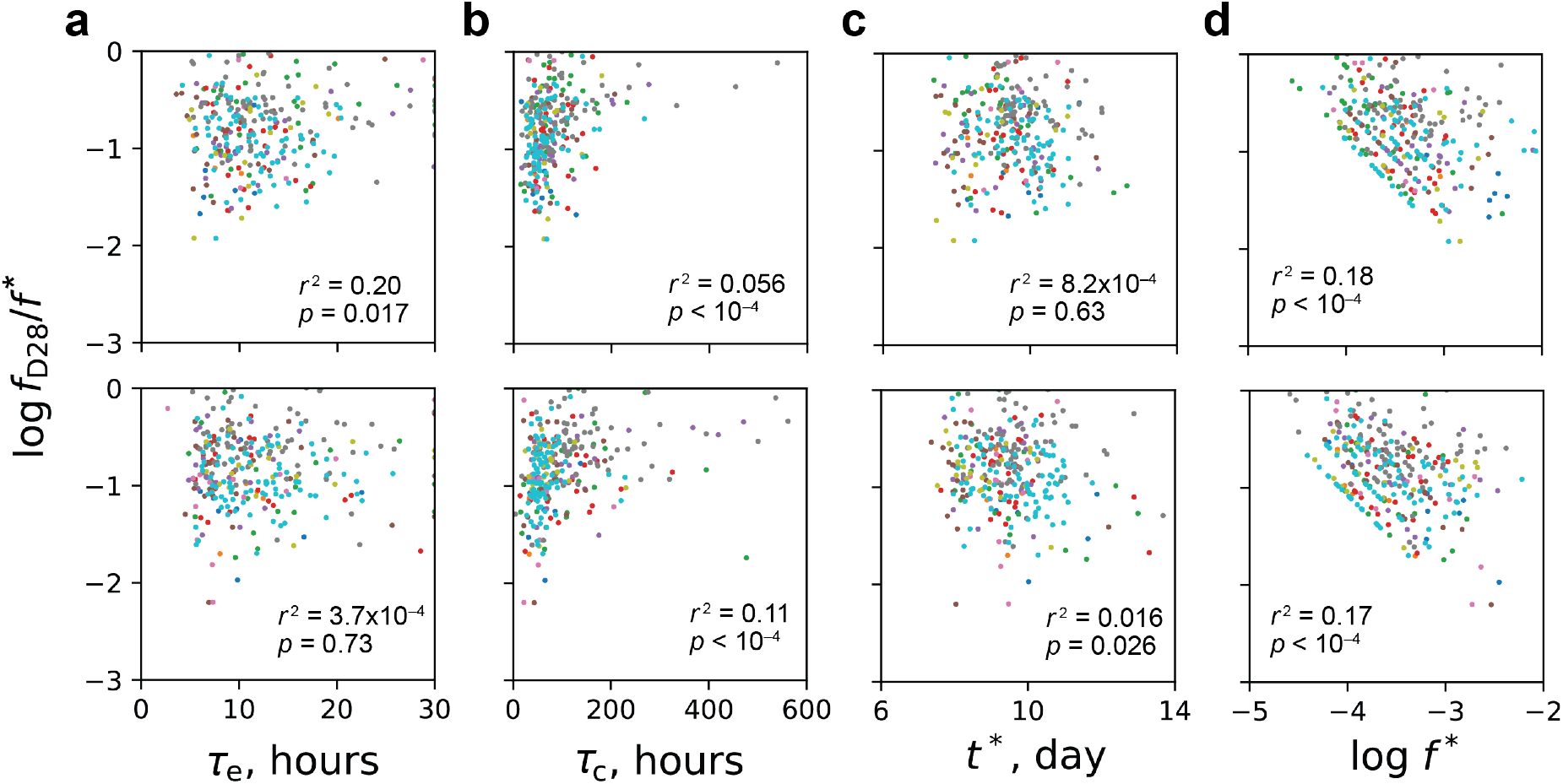
Correlation between clonal persistence and kinetic parameters. **a–d**, Relationship between the level of clonal persistence—quantified as log *f*_D28_*/f* ^∗^ and early kinetic parameters: clonal expansion timescale (**a**), contraction timescale (**b**), peak day (**c**), and peak abundance (d). Correlations were assessed using Pearson’s correlation test. Upper panels show measurements based on TCR *α*-chains; lower panels show *β*-chains. Different colors denote data from different individuals.

